# Pangenome of cultivated beet and crop wild relatives reveals parental relationships of a tetraploid wild beet

**DOI:** 10.1101/2023.06.28.546919

**Authors:** Katharina Sielemann, Nicola Schmidt, Jonas Guzik, Natalie Kalina, Boas Pucker, Prisca Viehöver, Sarah Breitenbach, Bernd Weisshaar, Tony Heitkam, Daniela Holtgräwe

## Abstract

Most crop plants, including sugar beet (*Beta vulgaris* subsp. *vulgaris*), suffer from domestication bottlenecks and low genetic diversity caused by extensive selection for few traits. However, crop wild relatives (CWRs) harbour useful traits relevant for crop improvement, including enhanced adaptation to biotic and abiotic stresses.

Especially polyploids are interesting from an evolutionary perspective as genes undergo reorganisation after the polyploidisation event. Through neo-and subfunctionalisation, novel functions emerge, which enable plants to cope with changing environments and extreme/harsh conditions. Particularly in the face of climate change, specific stress and pathogen resistances or tolerances gain importance. To introduce such traits into breeding material, CWRs have already been identified as an important source for sustainable breeding. The identification of genes underlying traits of interest is crucial for crop improvement.

For beets, the section *Corollinae* contains the tetraploid species *Beta corolliflora* (2n=4x=36) that harbours salt and frost tolerances as well as a wealth of pathogen resistances. The number of beneficial traits of *B. corolliflora* is increased compared to those of the known diploids in this section (all 2n=2x=18). Nevertheless, neither the parental relationships of *B. corolliflora* have been resolved, nor are genomic resources available to steer sustainable, genomics-informed breeding.

To benefit from the resources offered by polyploid beet wild relatives, we generated a comprehensive pangenome dataset including *B. corolliflora*, *Beta lomatogona*, and *Beta macrorhiza*, as well as a more distant wild beet *Patellifolia procumbens* (2n=2x=18). Joined analyses with publicly available genome sequences of two additional wild beets allowed the identification of genomic regions absent from cultivated beet, providing a sequence database harbouring traits relevant for future breeding endeavours. In addition, we present strong evidence for the parental relationship of the *B. corolliflora* wild beet as an autotetraploid emerging from *B. macrorhiza*.

## Background

### Sugar beet, crop wild relatives and the potential for breeding

The crop plant sugar beet (*Beta vulgaris* subsp. *vulgaris*) is of high economic relevance contributing to approximately 20% of the global sugar production (Biancardi and Lewellen 2020). To increase sugar production, early breeding focused mainly on yield. This domestication process introduced a strong genetic bottleneck resulting in diminished diversity available to breeders (Panella et al. 2020; Monteiro et al. 2018). Other important traits, like resistances to biotic and abiotic stresses, were initially neglected but gain more and more relevance in the face of climate change (Ristaino et al. 2021). It was already shown that some sea beets and some wild beets contain agronomically important traits that were lost during domestication (Biancardi and Lewellen 2020). Examples for such traits include salt and nematode tolerances (Panella et al. 2020; Cai et al. 1997; Capistrano-Gossmann et al. 2017). However, other crop wild relatives (CWRs) of sugar beet might harbour even more potential in terms of traits which can be incorporated to allow more sustainable beet cultivation (Panella et al. 2020). To this end, we sequenced and assembled the genomes of four different wild beets, namely *Beta corolliflora*, *Beta lomatogona*, *Beta macrorhiza*, and *Patellifolia procumbens*, for which no genome sequences were available until now.

### Pangenome instead of a single reference to identify ‘lost’ regions harbouring traits of interest

Several pangenome studies show that deep understanding of traits of interest requires the analysis of related species, whereas a single reference genome sequence often lacks important information, e.g. due to presence/absence variations (PAVs) between different species or accessions (Bayer et al. 2020, 2021). Since a pangenome of a taxonomic group is not static, we use the term pangenome synonymously with pangenome dataset, comprising sequencing reads, genome assemblies, and annotations of different related species. In the context of a crop pangenome study, the investigation of CWRs is of particular interest for breeding endeavours - not only to improve yield, but also to (re-)introduce regions lost during domestication which encode traits relevant for the defence against biotic and abiotic stresses. This is increasingly relevant due to climate change. Upcoming climatic conditions, including higher temperatures and heavy rain or flooding events, may promote favourable conditions for plant pests and diseases (Ristaino et al. 2021; Jabran et al. 2020). Therefore, we compared the genome sequences of sugar beet and CWRs to identify regions absent from the *B. vulgaris* subsp. *vulgaris* genome sequence but harbouring important trait-associated genes presumably relevant for the generation of enhanced varieties through breeding.

### Resolving the origin of the polyploid wild beet *Beta corolliflora*

The pangenome is not only of relevance at the gene or functional level, but also provides substantial insights into the evolution of crops and wild species (Bayer et al. 2020). Especially polyploid organisms are evolutionarily interesting as genes often undergo reorganisation and neo-or subfunctionalisation after the polyploidisation event. Novel functions can emerge, enabling the plant to better adapt to changing environments and stressful conditions (Adams and Wendel 2005; Otto and Whitton 2000; Van de Peer et al. 2017, 2021). This is not only true for allopolyploids, where the genomes of two different species are combined, but also for autopolyploids that evolve from one diploid parent. Despite the extensive niche overlaps of progenitor and descendant, these ploidy increases can stabilise heterosis, resulting for example in higher adaptability to stress (Van de Peer et al. 2021; Wang et al. 2013).

Regarding beets and wild beets, the section *Corollinae* harbours a range of higher polyploids. Among them, the most well-known is the tetraploid *Beta corolliflora* (2n=4x=36). It harbours a wide range of beneficial traits, including salt and frost tolerance as well as various resistances against pathogens (Panella et al. 2020). Yet, the type and origin of its polyploidy remain unclear. Having a diploid chromosome configuration of 2n=2x=18, *B. lomatogona* and *B. macrorhiza* are considered as potential parents. In contrast to *B. nana*, these species are the only known diploids of the section *Corollinae* that show a geographical distribution overlap with *B. corolliflora* (Sielemann et al. *B. corolliflora* is therefore considered to be either an allotetraploid resulting from hybridization of *B. lomatogona* and *B. macrorhiza*, or an autotetraploid resulting from a whole genome duplication of only one of those two species (or closely related to extinct relatives of one of those two species) (Frese and Ford-Lloyd 2020; Reamon-Büttner et al. 1996). A pangenome resource will be useful to trace the origin of *B. corolliflora*’s tetraploidy and may provide important insights into the past polyploidisation event.

### Objective

In this study, we present evidence for the tetraploid origin of *B. corolliflora* by generating the first genome sequence assemblies for four different sugar beet wild relatives - *B. corolliflora* (4x), *B. lomatogona* (2x), *B. macrorhiza* (2x), and as an outgroup *P. procumbens* (2x). These newly available beet genomic resources, together with the genome sequence of the cultivated sugar beet reference KWS2320 (assembly version KWS2320ONT v1.0; *B. vulgaris* subsp. *vulgaris*) (Sielemann et al. 2023), sea beet (*B. vulgaris* subsp. *maritima* WB42) (Rodríguez del Río et al. 2019), and *B. patula* (Rodríguez del Río et al. 2019), were used to gain insights into the beet pangenome by i) employing cytogenetic, *k-*mer-and gene-based methods to get evidence for the parental relationships of the tetraploid wild beet *B. corolliflora*, and by ii) identifying ‘lost’ regions in the cultivated sugar beet KWS2320 with relevance for breeding.

## Results

### Genome assemblies of wild beets

Three long read-based assemblies (*B. corolliflora*: BcorONT v1.0, *B. lomatogona*: BlomONT v1.0, and *P. procumbens*: PproONT v1.0) and a short read-based assembly (*B. macrorhiza*: Bmrh v1.0) of wild beet species were generated and serve as additional genomic resources for future analyses and breeding (Table 1).

**Table 1:** Assembly statistics of the new beet genomic resources. The assemblies of *B. corolliflora*, *B. lomatogona*, and *P. procumbens* are based on long reads, whereas the assembly of *B. macrorhiza* is based on short reads.

| Species | *B. corolliflora* | *B. lomatogona* | *P. procumbens* | *B. macrorhiza* |
| --- | --- | --- | --- | --- |
| Assembly name | BcorONT v1.0 | BlomONT v1.0 | PproONT v1.0 | Bmrh v1.0 |
| Data type | ONT | ONT | ONT | Illumina |
| Assembly size [bp] | 1963,172,020 | 1032,079,534 | 977,011,471 | 736,230,911 |
| Number of contigs | 4,355 | 1,530 | 1,542 | 218,216 |
| N50 [bp] | 720,692 | 1,746,059 | 1,392,274 | 7,104 |
| GC content [%] | 36.79 | 36.57 | 36.53 | 36.38 |
| Repeat content | 1,408,234,706 bp (71.73%) | 719,015,079 bp (69.67%) | 664,938,575 bp (68.06%) | 472,886,165 bp (64.23%) |
| BUSCOs<br>(n: 2326) | C:95.4% [S:22.5%,<br>D:72.9%], F:0.6%,<br>M:4.0% | C:92.8% [S:58.8%,<br>D:34.0%], F:0.6%,<br>M:6.6% | C:90.1% [S:43.8%,<br>D:46.3%], F:2.6%,<br>M:7.3% | C:68.0% [S:63.9%,<br>D:4.1%], F:12.3%,<br>M:19.7% |

The largest genome sequence assembly was constructed for the tetraploid *B. corolliflora* with a total size of approximately 1.96 Gb (Table 1). The genome sequence assemblies of the diploid species *B. lomatogona* and *P. procumbens* have a comparable approximate total assembly size of 1 Gb with 1500 contigs each. The final genome sequence assembly for *B. macrorhiza* comprises 218,216 contigs with a cumulative size of 736 Mb. Here, limited access to leaf material restricted DNA amounts, resulting in an Illumina-only assembly. All newly generated assemblies exceed the size of the KWS2320 sugar beet reference genome sequences Refbeet-1.2 and RefBeet-1.5 (Holtgräwe; Dohm et al. 2014; Minoche et al. 2015).

In general, all long read-based assemblies show high completeness with BUSCO percentages above 90%. The number of non-single copy (at least duplicated) BUSCOs is substantially higher for the tetraploid species (72.9%), with most of the complete BUSCOs being triplicated in BcorONT v1.0.

The genome assembly sequences of section *Corollinae* species (average of BcorONT v1.0, BlomONT v1.0, and Bmrh v1.0: 36.58%) show a higher GC content compared to section *Beta* (35.74% in KWS2320ONT v1.0). In addition, the genome assemblies of *Corollinae* species are substantially larger compared to species of the section *Beta* (see Table 1). The repeat content is similarly high in all genome assembly sequences ranging from 64.23% in Bmrh v1.0 to 71.73% in BcorONT v1.0, but generally higher in species with larger genomes.

### Resolving parental relationships of tetraploid *B. corolliflora*

To demonstrate the power of the wild beet genome sequencing and assemblies, we addressed the question regarding the parental relationships of tetraploid *B. corolliflora*. For this, we consider three different hypotheses that target the emergence from *B. lomatogona* and *B. macrorhiza* (Figure 1A). These hypotheses are: autotetraploidy originating from either diploid *B. lomatogona* (I) or *B. macrorhiza* (II) and allopolyploidy originating from hybridization of both diploid species (III).

**Figure 1:**
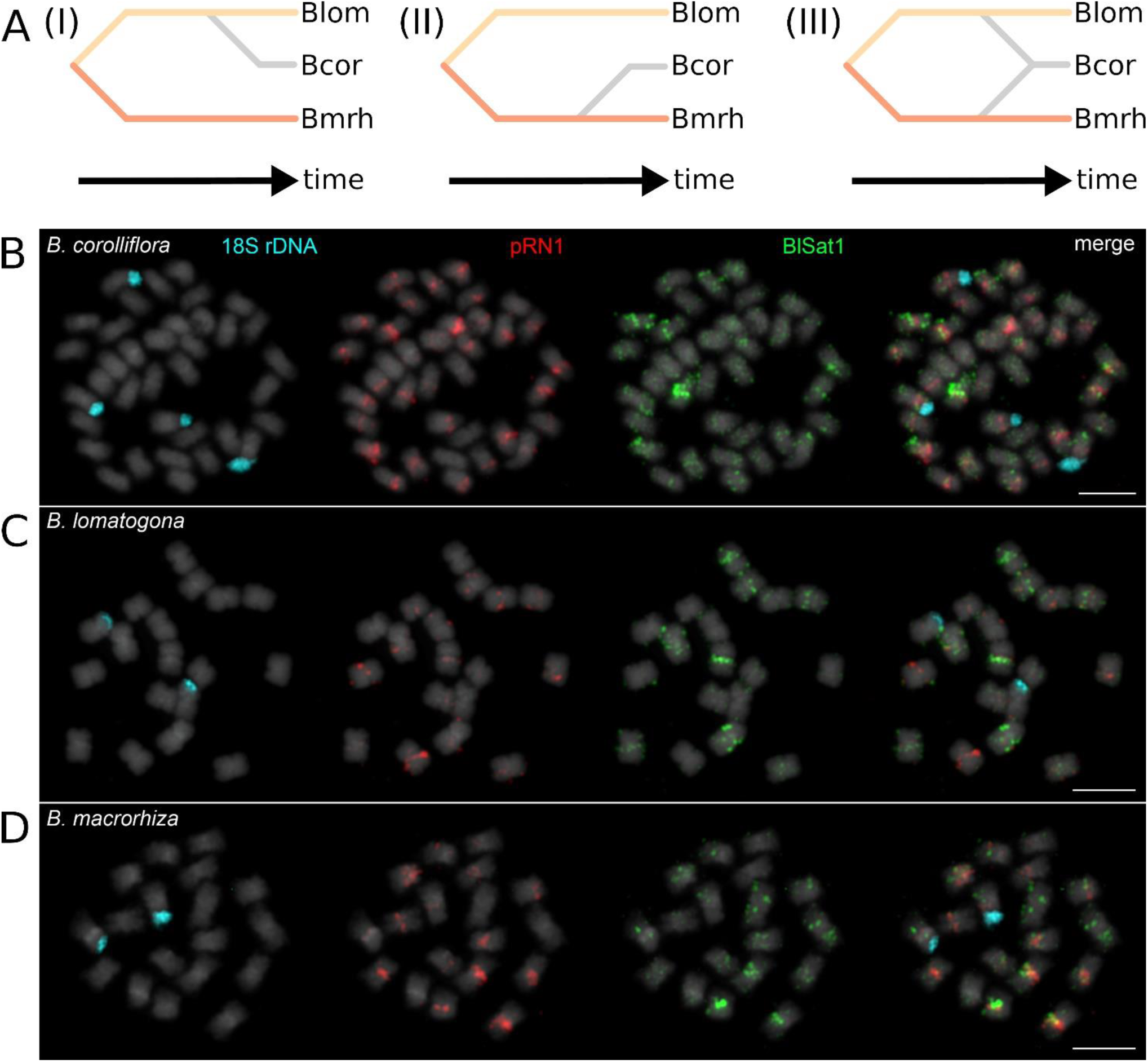
Possible parental relationships of tetraploid *B. corolliflora* (Bcor) and their support by cytogenetics. (A) Hypothesis (I) shows *B. corolliflora* as autotetraploid species with *B. lomatogona* (Blom) being the single parent species. Hypothesis (II) shows *B. macrorhiza* (Bmrh) as a single parent of autotetraploid *B. corolliflora* whereas hypothesis (III) considers both parents contributing to allopolyploid *B. corolliflora*. (B-D): Chromosomal landmarks along mitotic chromosomes of *B. corolliflora*, *B. lomatogona* and *B. macrorhiza* are not sufficient to unequivocally deduce the parental relationships. Mitotic chromosomes of the wild beets *B. corolliflora* (A), *B. lomatogona* (B), and *B. macrorhiza* (C) were hybridised with probes marking the 18S rDNA gene (with DY415; blue signals) and the satellite DNAs pRN1 (with streptavidin-Cy5; red signals), BlSat1 (with antidigoxygenin-FITC; green signals). The 18S rDNA is a widely used cytogenetic mark, usually flagging one chromosome pair in beets (Paesold et al. 2012; Rodríguez del Río et al. 2019).The chromosomes were counterstained with DAPI (grey). See Supplemental_File_S1 for signal counts and interpretation. The scale bars correspond to 5 µm. Cytogenetically, hypothesis (II) is most supported, but evidence is not yet conclusive.

To better illustrate this question, we first outline how all three genomes compare on a chromosomal level. This follows a simple rationale: depending on the mechanism of tetraploidization, the chromosomes from the diploids should be found again in the chromosomal set of the tetraploid. Here, we show a cytogenetics approach with three probes based on tandemly repeated DNAs (Figure 1; Supplemental_File_S1).

The 18S rDNA probe, a widely used cytogenetic mark (Figure 1B-D, blue), distinctly labels four chromosomes in the tetraploid (Figure 1B) and two chromosomes in the diploids (Figure 1C, 1D), supporting all three hypotheses. Therefore, as the remaining two probes, we chose tandemly repeated satellite DNAs that occur solely in wild beets of the *Corollinae* and have potential to inform about genetic differences between the wild beet species. For beetSat10-pRN1, we observe hybridization on 32 chromosomes, with many major and moderate signals in *B. corolliflora* (Figure 1B red; signal counts in Supplemental_File_S1). Similarly, beetSat8-BlSat1 hybridizes to all chromosomes with varying intensity (Figure 1B green; signal counts in Supplemental_File_S1). Then, we comparatively hybridized these probes to *B. lomatogona* and *B. macrorhiza* chromosomes (Figure 1C, 1D; Supplemental_File_S1, A) to deduce expected signal counts for each hypothesis and to calculate how each count varies from the expectation (Supplemental_File_S1, B-D). As a result, we find least variance between the observed and the expected signal counts for hypothesis (II). Hence, we conclude most cytogenetic support for *B. corolliflora*’s emergence through autotetraploidization of *B. macrorhiza*, but also acknowledge the limitations of the analysis.

To explore the power of the wild beet genome data and assemblies for deducing and verifying the tetraploid parentage of *B. corolliflora*, we deployed five different computational approaches. Some of these approaches are based directly on Illumina reads as input data (read-based approaches) and are therefore not dependent on any assembly quality parameters.

The similarity of the genome sequences of two species reflects the distance of their relationship. In turn, the similarity of two sequences is reflected by the similarity of their *k*-mer sets. Essentially, the set of *k*-mers of a sequence equals a compact representation of that sequence. Additionally, comparing *k*-mer sets is assumed to be more robust than directly comparing sequences considering assembly errors, e.g. at repetitive regions. The horizontal bar plot (Figure 2A) summarises the composition of the *B. corolliflora* 21-mer set. *B. corolliflora* shares 27% of its *k*-mers exclusively with *B. macrorhiza* and 7% exclusively with *B. lomatogona*.

**Figure 2:**
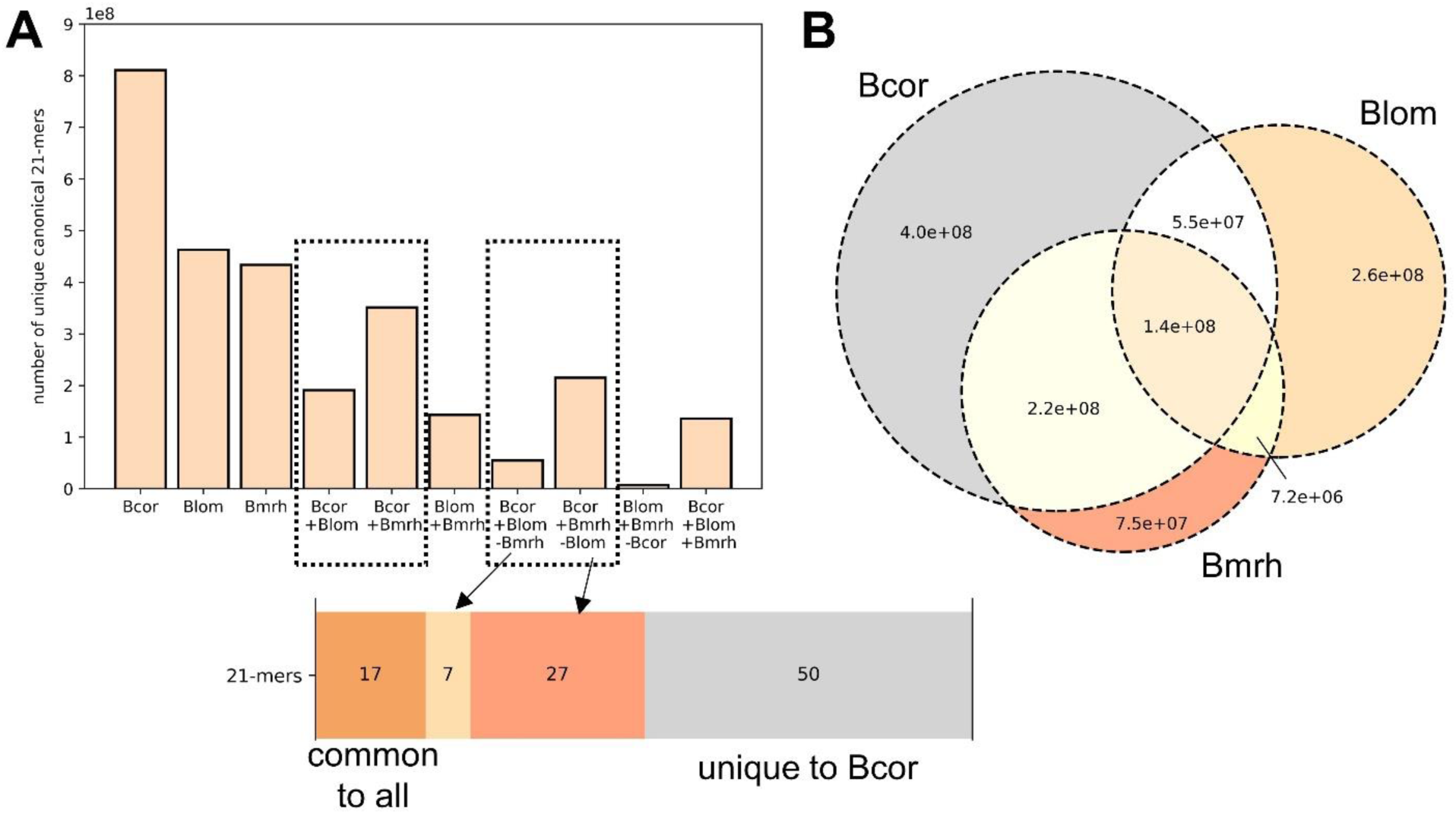
Results of the *k*-mer set operations for each tested hypothesis. A) Size (number of unique canonical 21-mers) of all investigated sets. The bars within the left black box represent intersections between the child species and each parent. The box on the right side represents the 21-mer set sizes including 21-mers present in only one candidate parent species but not in the other. The horizontal bar plot below summarises the composition of the *B. corolliflora* 21-mer set. B) Venn diagram for the read-based 21-mer sets of *B. corolliflora*, *B. lomatogona*, and *B. macrorhiza*.

The overlap of the read-based *k*-mer sets of *B. corolliflora*, *B. lomatogona*, and *B. macrorhiza* was visualised in a Venn diagram (Figure 2B). The *k*-mer set of *B. macrorhiza* has a substantially higher intersection/overlap with the *B. corolliflora k*-mer set (3.5e8; 81% of the whole *B. macrorhiza* set) compared to the *B. lomatogona k*-mer set (1.9e8; 41% of the whole *B. lomatogona* set). Comparable results were observed using assembly datasets instead of read datasets as input to generate the *k*-mer sets.

In a second approach, generalised trio binning was performed. We adapted the classical trio binning approach to resolve parental or more generally phylogenetic relationships by arguing that the number of reads assigned to one of the parent candidates reflects its relationship to the child relative to the other potential parent’s relationship to the child species. For both, the assembly-and the read-based trio, including *B. corolliflora*, *B. lomatogona*, and *B. macrorhiza*, the percentage of reads assigned to *B. macrorhiza* (40% and 51.9%) is substantially higher than the percentage of reads assigned to *B. lomatogona* (11.5% and 3.1%) (Table 2).

**Table 2:** Results of the generalised trio binning approach. For each trio, the type of analysis, the type of input datasets used to generate the *k*-mer sets, the name of the child species as well as the proportion of reads assigned to each of the four classes are provided. Abbreviations: Bcor = *B. corolliflora*, Blom = *B. lomatogona*, Bmrh = *B. macrorhiza*, Bnap = *B. napus*, Bole = *B. oleracea*, Brap = *B. rapa*, Bvul = *B. vulgaris* subsp. *vulgaris*, Bpat = *B. patula*, Bmar = *B. vulgaris* subsp. *maritima*.

| Analysis | Input | Child species | Parent A | Parent B | Unclassified | Chimeric |
| --- | --- | --- | --- | --- | --- | --- |
| Test case | Reads | Bcor | Blom: 0.031 | Bmrh: 0.519 | 0.173 | 0.029 |
| Test case | Assemblies | Bcor | Blom: 0.115 | Bmrh: 0.4 | 0.257 | 0.05 |
| Control allo | Reads | Bnap | Bole: 0.291 | Brap: 0.245 | 0.061 | 0.009 |
| Control ‘auto’ | Assemblies | Bvul | Bpat: 0.047 | Bmar: 0.356 | 0.307 | 0.066 |

As a control trio for a well-known allopolyploid species complex, datasets of *Brassica oleracea* and *Brassica rapa* as known parents of *Brassica napus* were analysed. A similar proportion of *B. napus* reads was assigned to both parental species (0.291 and 0.245). Normalising the results for the genome size differences (here, *B. napus* is considered both genomes combined (696+529)) results in a proportion of reads, assigned to *B. oleracea* (contributes 56.8% to the *B. napus* genome) and *B. rapa* (contributes 43.2% to the *B. napus* genome), of 0.512 and 0.567, respectively. The similar amount of *B. napus* reads assigned to both parents shows that the method leads to the expected results.

As an ‘autopolyploid’ control, *B. vulgaris* subsp. *maritima* as known progenitor of *B. vulgaris* subsp. *vulgaris* was used together with *B. patula* which is not considered to be a progenitor of *B. vulgaris* subsp. *vulgaris*. These species are no polyploids, however, the progenitor-descendant relationship of these species is known, which enables further validation of our approach. For this trio, a clear signal towards *B. vulgaris* subsp. *maritima* is visible (35.6%) whereas 4.7% of the reads are assigned to *B. patula*.

In a third *k*-mer based approach, *k*-mer fingerprints for numerous random sets were computed. As already described for the *k*-mer set operations approach, the more closely related two species are, the more similarity is expected between the respective *k*-mer sets.

The average absolute fingerprint sizes are shown in Table 3. *B. corolliflora* has the largest average fingerprint size (4,358). *B. patula*, *B. vulgaris* subsp. *maritima*, and *B. vulgaris* subsp. *vulgaris* show a similar average fingerprint size in the range of 3,061 to 3,088. *B. macrorhiza* shows the largest fingerprint intersection (3093; 92.7%), i.e. the overlap between the *B. macrorhiza* set with the set of the child species *B. corolliflora* (Table 3). This value is substantially higher than the one of the other putative diploid parent *B. lomatogona* (2822; 77.1%). Considering the diploids from the section *Beta*, *B. patula*, *B. vulgaris* subsp. *maritima*, and *B. vulgaris* subsp. *vulgaris* have similar fingerprint intersection sizes (2235-2251, 73.0%). The smallest fingerprint intersection size is observed for the wild beet representative from the sister genus *Patellifolia*, *P. procumbens* (2179; 66.4%).

**Table 3:**
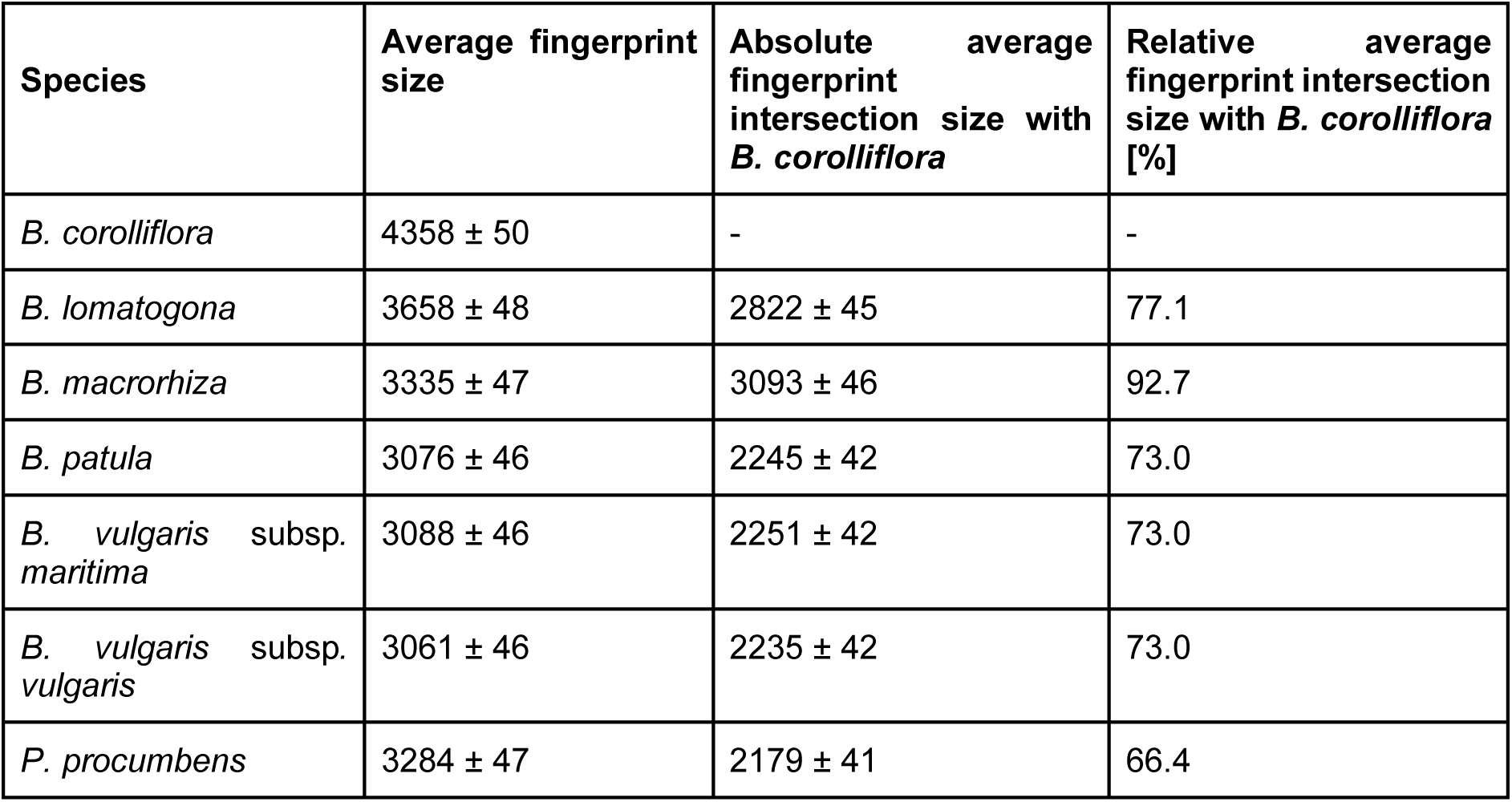
Results of the *k*-mer fingerprinting approach. 2^nd^ column: Average fingerprint sizes of all random sets of size 10,000 for all investigated species. 3^rd^ and 4^th^ column: Absolute and relative (relative with respect to the parent candidate) average fingerprint intersection sizes of all parent candidates. In addition to the average values, the standard deviations are shown.

| Species | Average fingerprint size | Absolute average fingerprint intersection size with *B. corolliflora* | Relative average fingerprint intersection size with *B. corolliflora* [%] |
| --- | --- | --- | --- |
| *B. corolliflora* | 4358 ± 50 | - | - |
| *B. lomatogona* | 3658 ± 48 | 2822 ± 45 | 77.1 |
| *B. macrorhiza* | 3335 ± 47 | 3093 ± 46 | 92.7 |
| *B. patula* | 3076 ± 46 | 2245 ± 42 | 73.0 |
| *B. vulgaris* subsp. *maritima* | 3088 ± 46 | 2251 ± 42 | 73.0 |
| *B. vulgaris* subsp. *vulgaris* | 3061 ± 46 | 2235 ± 42 | 73.0 |
| *P. procumbens* | 3284 ± 47 | 2179 ± 41 | 66.4 |

The fourth applied approach relied on cross-species mapping of synthetic reads. It is expected that the closer two species are related, the higher their sequence similarity. Therefore, it is expected to find more regions of an assembly of one species in an assembly of the other species, the closer these species are related. Based on these assumptions, a mapping approach was developed to present evidence for the parental relationships of tetraploid *B. corolliflora*. This mapping approach to resolve the parental relationships of *B. corolliflora* directly compares the two candidate parental species. Supplemental_File_S2 shows the percentage of synthetic *B. corolliflora* reads that mapped exclusively to *B. lomatogona*, exclusively to *B. macrorhiza* or to both species with a sequence identity of at least 60%. For all considered synthetic read lengths (5 kb, 10 kb, and 20 kb), the percentage of reads that map to both potential parents is below 1%. With shorter read length, the percentage of reads mapping only to *B. macrorhiza* increases whereas the percentage of reads mapping only to *B. lomatogona* decreases. For 5 kb reads, more than twice as many successfully mapped reads map exclusively to Bmrh v1.0 (68% versus 31% for BlomONT v1.0). When mapping 5 kb reads against the reference consisting of *B. lomatogona* shredded into 5 kb chunks and the short read assembly of *B. macrorhiza*, the results are almost identical to the ones using the full-length *B. lomatogona* assembly as reference.

In a fifth approach, that is based on gene sequences, the similarity of orthologous genes was used as a measure to assess the putative parents of tetraploid *B. corolliflora*. A large basis of single nucleotide variants (SNVs) in single-copy BUSCO genes was employed to calculate phylogenetic distances of the gene sequences of the potential parents to the respective gene sequence of *B. corolliflora*. For this, 140 single BUSCO gene phylogenies were computed. The phylogenetic distance of the *B. macrorhiza* genes to the respective closest related *B. corolliflora* gene (mean approx. 0.0213) is significantly smaller when compared to *B. lomatogona* (mean approx. 0.0305) (U-test; p≈5e-24) (Figure 3A). This means that *B. macrorhiza* genes are substantially more often found in a common phylogenetic unit together with the respective *B. corolliflora* gene, whereas *B. lomatogona* genes are often found on a separate branch in the phylogenetic tree (Figure 3B, 3C).

**Figure 3:**
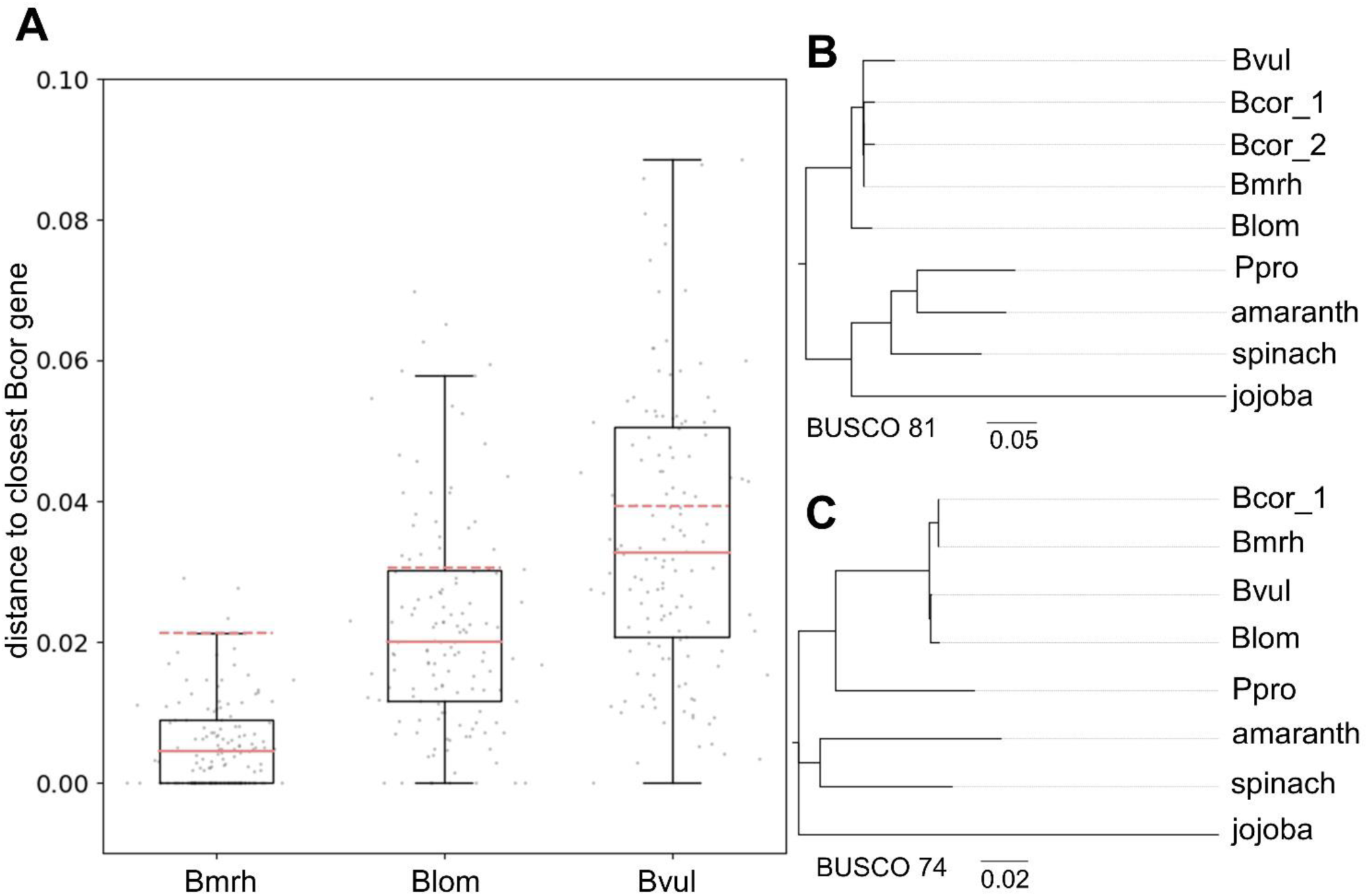
Results of the gene-based approach to determine the parental relationships of *B. corolliflora*. A) Phylogenetic distance of all ‘parental’ genes to the respective closest related *B. corolliflora* gene. The mean is shown as a dashed orange line whereas the median is represented by a solid orange line. B, C) Phylogenetic ML trees for two selected BUSCO genes. Bcor_1 and Bcor_2 represent two different copies of the same gene in *B.corolliflora* (duplicated BUSCO). Spinach (*S. oleracea*), amaranth (*A. hypochondriacus*), and jojoba (*S. chinensis*) were used as outgroup species. Abbreviations: Bcor = *B. corolliflora*, Blom = *B. lomatogona*, Bmrh = *B. macrorhiza*, Bvul = *B. vulgaris* subsp. *vulgaris*, Ppro = *P. procumbens*.

To gain more insight, phylogenetic distances in trees in which the *B. macrorhiza* gene is not clustered with the closest *B. corolliflora* gene were further investigated. Branch lengths in such trees are particularly small and *B. corolliflora*, *B. lomatogona*, and *B. macrorhiza* sequences of the respective genes are hardly distinguishable with phylogenetic distances of e.g., < 0.0086 (average distance between two sequences in this tree: 0.104197).

### ‘Lost’ regions in sugar beet but present in the wild beets

As especially polyploid CWRs might harbour properties/traits not present in the cultivated beet, the newly generated pangenome resources, including tetraploid *B. corolliflora*, were used to identify regions not present in the KWS2320 sugar beet breeding material. These regions might harbour information for traits relevant for breeding which are not present in the cultivated beet. Based on overlapping genes associated with specific traits, these regions are possibly interesting for future breeding endeavours. For the investigated CWRs, 4.0% to 10.2% of the genome sequence assemblies were found to be ‘zero coverage regions’ and therefore to be ‘lost’ and/or not present in sugar beet KWS2320 (Supplemental_File_S3). In these regions of the CWRs, several genes related to plant defence, to pathogens, to response to various stimuli or to other possibly interesting traits were identified.

## Discussion

To investigate the pangenome of sugar beet and its CWRs, the first genome sequence assemblies for four different wild beets (*B. corolliflora* (2n=4x), *B. lomatogona* (2n=2x), *B. macrorhiza* (2n=2x), and *P. procumbens* (2n=2x)) were generated. Published genome sequences of *B. patula* and *B. vulgaris* subsp. *maritima* (Rodríguez del Río et al. 2019) as well as the reference genome sequence of cultivated sugar beet (KWS2320ONT v1.0) (Sielemann et al. 2023), were integrated to i) get evidence for the parental relationships of the tetraploid beet *B. corolliflora* and ii) analyse genomic regions in CWRs associated with traits of interest.

### *B. macrorhiza* as single parent of autotetraploid *B. corolliflora*

We combined multi-colour cytogenetics with five computational approaches to elucidate the type of tetraploidy in *B. corolliflora* and its ancestry. Using all six approaches, we can now confidently exclude *B. lomatogona* as parental species, and we find comprehensive evidence of an emergence as autotetraploid from *B. macrorhiza*. Alternatively, as an option that we cannot distinguish from the autotetraploid scenario, *B. corolliflora* might be an allotetraploid derived from to different but closely related *B. macrorhiza* genotypes.

To define the diploid ancestry of a polyploid is a question that is and has been commonly addressed using cytogenetics (Schmidt et al. 2019; Heitkam et al. 2020; Desel 2002). Here, as the parental genomes are closely related with relatively limited variation amongst cytogenetic probes, the question is answered only with difficulty and not conclusively. Still, our cytogenetic analysis retained most support for *B. corolliflora*’s autotetraploidy emerging from *B. macrorhiza*.

To convincingly resolve the question of *B. corolliflora*’s tetraploidy, we leveraged five data-driven genomics approaches using our wild beet pangenome dataset. Three approaches are based on *k*-mers, one is based on mapping of synthetic reads and a fifth approach is based on sequences of conserved and orthologous genes. The advantage of the *k*-mer approaches using reads as input to generate the species-specific sets is that these approaches do not rely on a reference genome sequence and are not dependent on e.g. the identification of homology through computationally expensive (whole genome) alignment approaches (Ondov et al. 2016; VanWallendael and Alvarez 2022).

For all *k*-mer based approaches, it is important to take the genome size of the potential parents into account. The *k*-mer set size is dependent on the genome size and also on the size of the assembly, since the probability of a *k*-mer occurring just by chance grows with increasing genome sequence size. *B. corolliflora* has an assembly size of 1,963 Mb and a 21-mer set size of 6.7e8, whereas *B. lomatogona* and *B. macrorhiza* have an assembly size of 1,032 Mb and 736 Mb and a corresponding 21-mer set size of 4.9e8 and 4.3e8, respectively (see Table 1). For polyploids, the haploid genome size might be more relevant. An additional copy of a genome, e.g. diploid vs. autotetraploid, does not increase the *k*-mer set size linearly. However, rearrangements, TE expansions, and particularly small mutations occurring after the polyploidisation/hybridisation increase the potential for additional *k*- mers. In addition to the biological genome size, the completeness and therefore the quality of the assembly has similar effects on the *k*-mer set size. Even though the assembly quality for *B. macrorhiza* is lower compared to the quality of the long read assemblies, there is a striking signal towards *B. macrorhiza* for all approaches.

The composition of the *B. corolliflora* 21-mer set (Figure 2A) shows a higher overlap with the *B. macrorhiza* set than with the *B. lomatogona* set, indicating a closer relationship of *B. corolliflora* and *B. macrorhiza*. A higher *k*-mer set similarity implies a higher sequence similarity and therefore closer phylogenetic relationship. As visualised in the Venn diagram (Figure 2B), both the absolute and the relative intersection sizes of *B. corolliflora* and *B. macrorhiza* are greater than those of *B. corolliflora* and *B. lomatogona*, i.e. the 21-mer sets of *B. corolliflora* and *B. macrorhiza* are more similar. The Venn diagrams of the 21-mer sets of *B. corolliflora*, *B. lomatogona*, and *B. macrorhiza* based on assemblies and reads, respectively, are comparable. Especially considering the fragmented assembly of *B. macrorhiza*, this supports the robustness of the *k*-mer set operations.

For trio binning, if both investigated species were the actual parental species of the child, it was expected that approximately the same number of reads would be assigned to both supposed parents. If only one of the candidate species was the parent, substantially more reads should be assigned to the designated species. This number depends on the phylogenetic relationship of the second candidate to the child species. Multiple factors, however, may lead to a divergence from these expectations: unequal genome sizes of both parents lead to a higher expected number of reads assigned to the species with the larger genome. Bias during the process of sequencing may also lead to an uneven distribution of reads (Ross et al. 2013). Rearrangements and sequence differences originating during the species’ evolution, especially of the genome of the child species may distort read distribution and *k*-mer content. Since rearrangements and extended genome divergence are regularly observed in polyploid species (Van de Peer et al. 2017), the trio binning approach is mainly aimed at resolving the parental relationships of young hybrid species.

Two ‘control trios’ were selected to validate the generalised trio binning approach. As allotetraploid control, the *B. napus*, *B. oleracea*, and *B. rapa* trio was used (Lu et al. 2019). A similar number of reads was assigned to both known parents of *B. napus*. The number is slightly higher for *B. oleracea*, which can be explained by the larger 21-mer set size (2.3e8 for *B. oleracea* vs. 1.6e8 for *B. rapa*). Overall, the results show that the method leads to the expected results. The second control trio comprises the sea beet as known progenitor of sugar beet (Biancardi and Lewellen 2020; Wascher et al. 2022) as well as *B. patula*, not a progenitor of sugar beet. More than seven times more reads are assigned to the sea beet compared to *B. patula*, which confirms the close relation of sea beet and sugar beet.

Regardless of using assemblies or reads as input for a trio of interest, substantially more reads are assigned to *B. macrorhiza*. Again, an advantage of this approach is that it does not rely on assembled data, even though it is possible to use assembled data as input. Using assemblies as input, *B. macrorhiza* obtains about four times more reads, whereas using reads as input, about 17 times more reads are assigned to *B. macrorhiza* than to *B. lomatogona*. These results indicate that *B. macrorhiza* might be the single parent of autoploid *B. corolliflora*. The difference in the results when using assemblies versus reads as input can be explained by the *k*-mer set sizes derived from the assemblies of *B. lomatogona* (3.5e8) and *B. macrorhiza* (2.8e8). The assembly-based *k*-mer set for *B. macrorhiza* is smaller, which can be explained by the fragmented short read assembly in which *k*-mers exclusive to unassembled regions might be missing. Using reads as input, it can be assumed that the normalised read datasets reflect the true *k*-mer sets well. Indeed, the difference in the size of the exclusive *k*-mer sets when using reads is smaller (3.2e8 for *B. lomatogona* and 2.9e8 for *B. macrorhiza*).

The idea of the *k*-mer fingerprinting approach is similar to the *k*-mer set operations method, however, there are two major differences: i) the randomisation introduced in the fingerprinting method can reduce the impact of errors when taking the average over a sufficiently large number of random sets and ii) this approach allows to compare more than two parent candidates simultaneously. The average fingerprint sizes are mainly related to genome size and sequence diversity (Table 3). *B. corolliflora* shows the largest average fingerprint size since the genome sequence is the largest among the investigated organisms. Considering the relative fingerprint intersection sizes, the results reflect the phylogenetic relationships of the species (Sielemann et al. 2022). *P. procumbens* has the highest phylogenetic distance to *B. corolliflora* among the investigated organisms and shows the smallest relative fingerprint intersection size. The fingerprint intersection sizes of all other investigated species also directly reflect the phylogenetic distances. The substantial difference in average relative fingerprint intersection size between *B. lomatogona* (77.1%) and *B. macrorhiza* (92.7%) suggests that *B. macrorhiza* is more closely related to *B. corolliflora* and presumably the single parent species. For the *k*-mer fingerprinting approach, an additional comparison with *B. nana* and *B. intermedia*, two additional species of the section *Corollinae*, would have been interesting, however, not enough data was available.

The synthetic read mapping approach reflects the similarity between sequence sections (synthetic reads) of *B. corolliflora* and the assembly sequences of the potential parent species. An advantage of this approach is the equal coverage distribution of the child species’ synthetic reads close to one. Therefore, specific regions are not substantially over-or underrepresented and the results are not biased by such sequences. Further, such synthetic, contig-based reads likely contain fewer errors than the actual sequencing reads the contigs are based on. The decrease in the percentage of reads which exclusively map to *B. macrorhiza* with increasing synthetic read length (Supplemental_File_S2), can be explained by the high fragmentation of the *B. macrorhiza* assembly. For synthetic reads of 5 kb length, more than twice as many reads map exclusively to *B. macrorhiza* compared to *B. lomatogona*. This indicates a higher sequence similarity and thus also a closer relationship between *B. macrorhiza* and *B. corolliflora* as opposed to *B. lomatogona* and *B. corolliflora*.

The gene-based approach was developed to assess the sequence similarity of conserved BUSCO genes (Simão, Felipe A and Waterhouse, Robert M and Ioannidis, Panagiotis and Kriventseva, Evgenia V and Zdobnov, Evgeny M 2015) between child and potential parent species. These investigated sequences are more similar between *B. macrorhiza* and *B. corolliflora* as shown by the clusters in the phylogenetic tree separate from the respective *B. lomatogona* gene sequence.

Combining all our results, cytogenetics and the five computational approaches, a clear pattern towards resolving *B. corolliflora*’s ancestry emerges (Table 4).

**Table 4:**
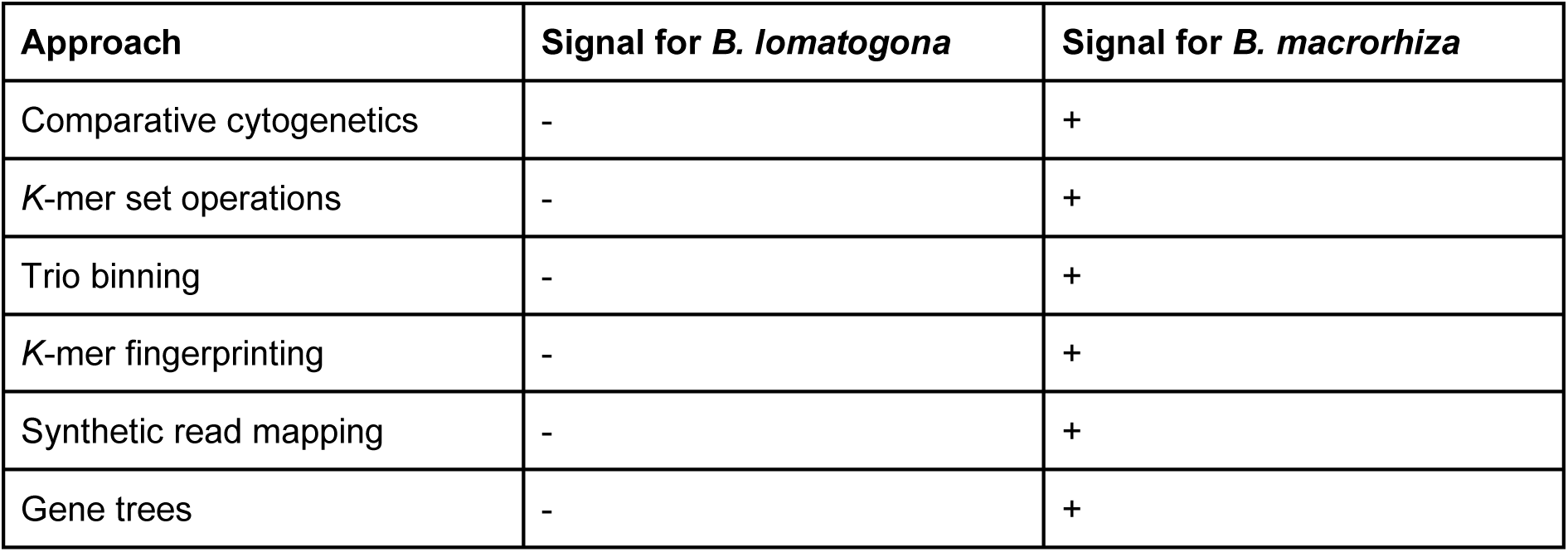
Overview of the results of each of the five newly developed methods to get evidence for the parental relationships of *B. corolliflora*. A plus (+) indicates that the method yields a signal for the corresponding species, while a hyphen (-) means that the respective species is not likely to be in a parental relationship with the tetraploid wild beet *B. corolliflora*.

All approaches show a clear signal towards *B. macrorhiza* being the single parent species of *B. corolliflora*, which therefore would be most likely an autotetraploid species. However, we cannot exclude the possibility that the real parental species of *B. corolliflora* is an unknown and possibly already extinct species very closely related to *B. macrorhiza*, or that *B. corolliflora* originated from a hybridisation event of such an unknown species with *B. macrorhiza* (Figure 4). Further, allo-and autopolyploidy are considered to reside along a ‘spectrum’ (Mason and Wendel 2020): i) highly diverse subgenomes from a single species can lead to the formation of a more polymorphic autopolyploid as compared to an allopolyploid species derived from two less diverged species. ii) Homoeologous exchanges can contribute to the formation of a partially autopolyploid species from an initial allopolyploid. This means that different regions of the genome appear to be allopolyploid whereas other regions appear to be autopolyploid. iii) Directional selection of genes, which favours one of the parental genomes, may cause homoelogs to ‘appear’ autopolyploid (Mason and Wendel 2020) even though the species is an allopolyploid of origin. Recreation of polyploids by crossing of the parental species may resolve if some of these mechanisms occur after the polyploidisation event.

**Figure 4:**
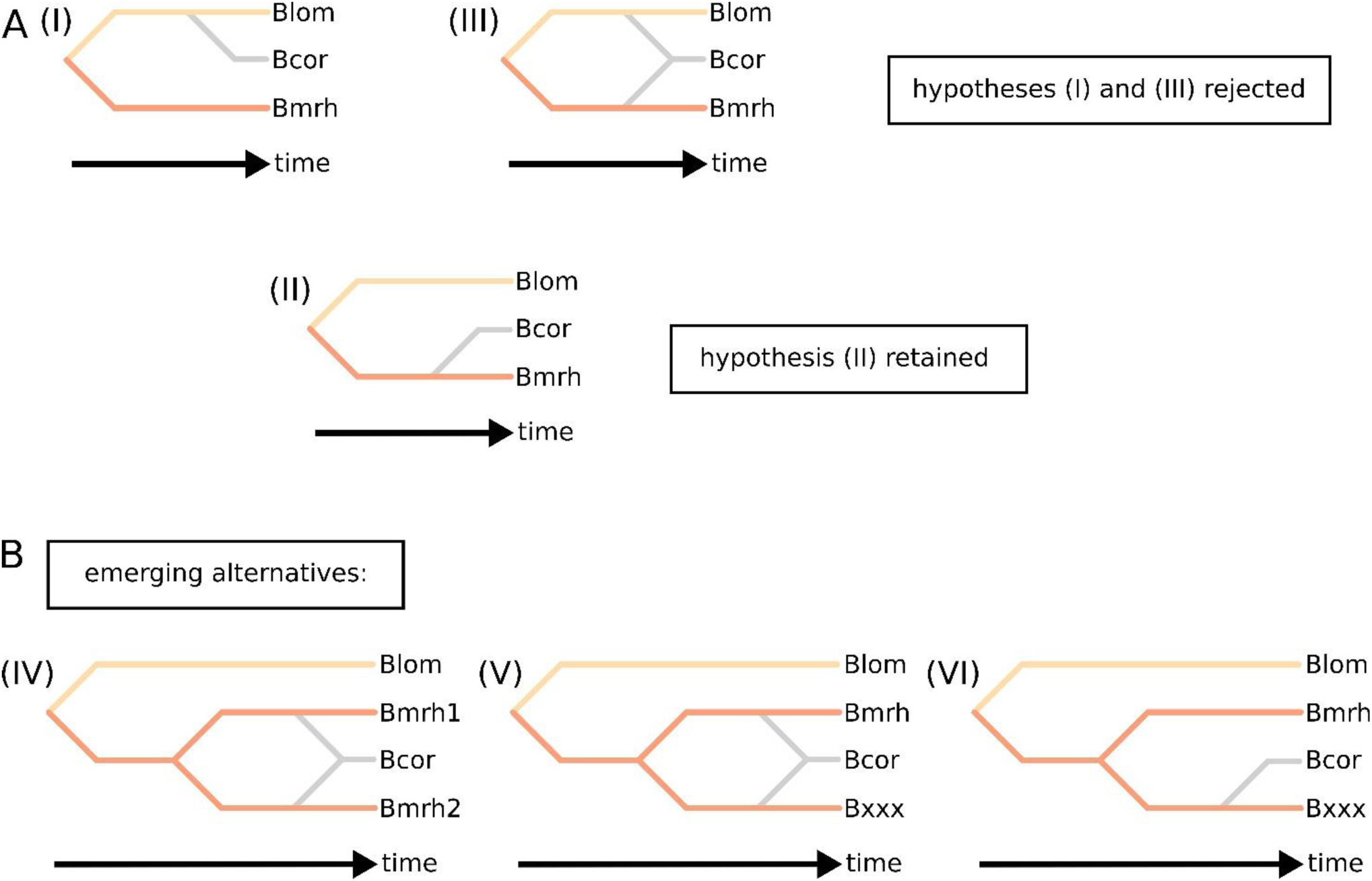
Amended hypotheses regarding the origin of tetraploid *B. corolliflora*. From the initial three hypothesis (A I-III), two hypotheses were disproved (I and III). *B. corolliflora* (Bcor) might be an autotetraploid species with Bmrh as a single parent (hypothesis II). This possibility is sharpened by the emergence of three new hypotheses (B IV-VI), in which *B. corolliflora* either originated from the hybridisation of two different *B. macrorhiza* cytotypes (IV), of *B. macrorhiza* with an unknown, possibly extinct *Beta* species (Bxxx) closely related to *B. macrorhiza*, or from the autopolyploidisation from this unknown *Beta* species (VI). However, these hypotheses cannot be tested with the available data as the existence of Bxxx is unknown.

### Harnessing CWRs to identify traits relevant for crop improvement

The pangenome dataset was used to identify CWR regions that are not present in cultivated sugar beet represented by KWS2320. ‘Lost’ regions in the cultivated sugar beet KWS2320, but present in the wild beet species, were defined as follows. If the region is not present in sugar beet, (almost) no reads of sugar beet should map to the corresponding region in any of the crop wild relatives. Based on this rationale, ‘zero coverage regions’ were extracted. Genes overlapping with ‘zero coverage regions’ were extracted since these regions might be relevant for future breeding endeavours. Potentially beneficial biotic and abiotic traits among the ’lost’ genes were identified. In the following, the detected traits are discussed in terms of their relevance for sugar beet breeding.

Through a functional annotation that was generated for BcorONT v1.0, BlomONT v1.0, Bmrh v1.0, and PproONT v1.0 (10.4119/unibi/2966932), the set of identified CWR genes was investigated for specific disease resistance genes, genes conferring tolerances, and genes related with response to bacteria, viruses, or fungi. Most genes identified in the zero coverage regions have no functional annotation, however, 39 different disease resistance proteins and putative disease resistance proteins were collectively identified for all four species in the functional annotations. Only 16 of them were present in the annotation of at least two species, the remaining 23 were unique to one of them. The annotation for *B. corolliflora* contained all of the five (putative) R-genes *RGA1-5*, *B. lomatogona* and *B. macrorhiza RGA1-4*, and *P. procumbens RGA3*. A *RGA2* homolog confers resistance to the oomycete *Phytophthora infestans* in wild potato (Song et al. 2003; van der Vossen et al. 2003). Additionally, multiple putative disease resistance proteins with no further known function were found. For *B. corolliflora* and *B. lomatogona*, the gene *RPP8* was found, which confers resistance to *Prenospora paraistica*, which is an oomycete (Berardini et al. 2015; Cooley et al. 2000; Zhu et al. 2011, 101). Multiple genes related to *A. thaliana* R-genes were found, one of which (*At3g14460/LRRAC1*, found in *B. corolliflora* and *B. macrorhiza*) is associated with defence response to fungal pathogens (Bianchet et al. 2019; Bairoch and Boeckmann 1991). *B. corolliflora* contained most unique resistance genes (22) compared to the other investigated species, followed by *B. macrorhiza* (20). Genes that are unique to *B. corolliflora* were e.g. *RPM1*, *At5g66890, At5g43730*, and the putative late blight resistance protein homolog *R1B-19* (*Solanum demissum*). *RPM1* confers resistance to some *Pseudomonas syringae* strains (Berardini et al. 2015; Yoon et al. 2022).

The *A. thaliana* orthologs (RBHs) were used for the transfer of functional information, especially for the two species (*B. patula* and *B. vulgaris* subsp. *maritima*) for which no other functional annotation was available. Several genes play a role in thermotolerance (e.g. response to heat/cold) and in response to various bacterial, viral and fungal pathogens. Further, genes associated with stress response to salt and drought, as well as genes associated with the regulation of flowering time, were identified. Genes relevant in response to herbivores include *KTI1* (*B. patula*) and *KTI5* (*B. macrorhiza*), which are involved in the defence response to spider mites (*Tetranychus urticae*) (Arnaiz et al. 2018). Spider mites infect a wide range of hosts, one of them being sugar beet (Reynolds et al. 1967). It has been shown that spider mites have a high amount of pesticide resistances, which is why a plants’ natural defence against them is beneficial (Arnaiz et al. 2018). Additionally, genes involved in the defence or response to some fungi, viruses (e.g. geminiviruses (Chung and Sunter 2014)), bacteria, as well as the nematode *Heterodera schachtii* (Shah et al. 2017) were identified. In the context of abiotic stresses, multiple genes relevant to drought resistance and tolerance of water deprivation were found (e.g. *B. macrorhiza*, *B. vulgaris* subsp. *maritima*). Due to climate change, drought displays a major limiting factor when it comes to sugar beet breeding (Ober and Rajabi 2010). Drought already causes about 10% of yield loss in parts of Europe and is believed to aggravate even further (Ober and Rajabi 2010). Another important abiotic factor is temperature. Genes related to heat acclimation (e.g. *B. corolliflora*), as well as cold response (e.g. *B. macrorhiza*) were identified. Sugar beets are predominantly cultivated in the temperate zone and grow most effectively in temperature ranges between 17 °C and 25 °C (Ober and Rajabi 2010). However, hotter and colder climates present potential new cultivation areas for adapted sugar beets. For example, freezing temperatures are harmful for sugar beet seedlings, which is why prior breeding initiatives already bred for cold resistant variants (Burenin et al. 1994). The findings suggest that wild beets might have a relevant potential to improve the adaptation of sugar beet to extreme climate conditions.

In terms of pathogen resistances, various examples of different categories could be identified. *B. vulgaris* subsp. *maritima* further contained a homolog (*At4g13350/NIG*) that negatively affects the tolerance against geminiviruses, a broad group of plant viruses. One of the viruses contained in that group is the beet curly top virus, which infects sugar beet (Yazdi et al. 2008). It causes curly top disease, which results in leaf curling, phloem necrosis and other symptoms (Yazdi et al. 2008). An important pathogen that has already been relevant in prior breeding initiatives is the cyst nematode *Heterodera schachtii*. A nematode resistance has been successfully transferred from *P. procumbens* to sugar beet in the past (Cai et al. 1997). In *P. procumbens* and *B. macrorhiza*, a gene (*At2g01340/At17.1*) which is associated with response to nematode infection, was identified. For *B. lomatogona*, *AT5G06860/PGIP1* was identified and this homolog attenuates infection with *Heterodera schachtii* (Shah et al. 2017). The gene *At2g01340/At17.1* is significantly induced in response to *Sclerotinia sclerotiorum* (pathogenic fungus), *Botrytis cinerea*, *Pseudomonas syringae* pv. *tomato* DC3000 *AvrRPS4*, *Verticillium dahliae*, and *Colletotrichum tofieldiae* (Didelon et al. 2020). A gene (*At2g43710/SSI2*) that is related to the response to the green peach aphid has been identified in *B. macrorhiza* (Berardini et al. 2015; Li et al. 2021). This insect has been shown to be an important transmitter of the previously mentioned curly top virus in sugar beet(Sylvester 1956). Mutation of this gene in *A. thaliana* causes hyper-resistance (Berardini et al. 2015; Li et al. 2021).

In summary, the presented method led to the identification of various regions and genes of interest. Even though the model organism *A. thaliana*, instead of sugar beet itself, had to be used to extract possible functions, the results show that the genetic variation present in beet wild relatives provides high potential to expand the sugar beet’s gene pool.

In a second approach, we identified regions derived from *B. vulgaris* subsp. *maritima* - the progenitor of sugar beet (Biancardi and Lewellen 2020) - and show evidence to support the assumption of sea beet being the progenitor of cultivated sugar beet (Supplemental_File_S4). Despite the usage of relatively strict thresholds to ensure the identification of high-confidence regions derived from sea beet, more than 101 Mb of highly conserved regions, representing 17.6% of the whole genome sequence, were identified in sugar beet. Conserved genes within these regions, possibly derived from sea beet, are e.g. associated with response to salt stress (more than 50 genes). Cultivated beets show higher salt tolerance compared to other crops, especially during germination and seed development (Pinheiro et al. 2018; Skorupa et al. 2019). The ability to tolerate high salt concentrations is a great advantage for wild sea beets since they are almost exclusively found in coastal regions (Romeiras et al. 2016). In such environments, salt stress represents the most significant abiotic stress. The results of our analysis and the mentioned studies suggest that many of the identified salt stress-related genes have originated in the sea beet.

## Conclusion

In this study, the pangenome dataset of sugar beet and CWRs was harnessed to get evidence for the parental relationships of a polyploid species and to identify traits relevant for crop improvement.

The developed methods to resolve polyploid relationships are based on different concepts and lead to unambiguous results concerning the three tested hypotheses. Therefore, *B. lomatogona* can be excluded as parent species of *B. corolliflora.* Further, it can be concluded that *B. macrorhiza* might be the single parent of the autotetraploid wild beet *B. corolliflora*. The newly developed approaches used to solve this question can also be applied to other datasets. The generalised trio binning approach seems promising to resolve parental relationships and in general phylogenetic relations of closely related species. Extending the generalised trio binning approach to not only consider unique *k*-mers but also *k*-mer frequencies is another interesting option. Further, all *k*-mer based methods could be used with skip-mers instead, a concept to include information from more distant genomic positions and to decrease the impact of SNVs (Clavijo et al. 2017).

The investigation of genomic regions not (anymore/yet) present in the cultivated sugar beet genome revealed several genes associated with pathogen resistance and tolerance to abiotic stresses. These genes are candidates for breeding endeavours to obtain sustainable crops.

Summarizing, we show the potential of the newly generated genome resources of CWRs of sugar beet which are an essential building block for future investigations and crop improvement.

## Methods

### Plant material

The Leibniz Institute of Plant Genetics and Crop Plant Research Gatersleben (IPK), Germany, provided seeds for *B. corolliflora* (BETA 408), *B. lomatogona* (BETA 674), *B. macrorhiza* (BETA 830), and *P. procumbens* (BETA 419). The material was transferred under the regulations of the standard material transfer agreement (SMTA) of the International Treaty. All plants were grown under standard greenhouse conditions.

### DNA extraction, sequencing and *de novo* assembly

For *B. macrorhiza*, a short read Illumina assembly was generated, as only a low amount of plant material was available, which was not sufficient for preparing DNA suitable for long read sequencing. High molecular weight DNA was extracted using a previously described CTAB-based method (Siadjeu et al. 2020). DNA extraction as well as Illumina sequencing was performed as previously described (Sielemann et al. 2022). In total, 138 GB read data were generated (Supplemental_File_S5). The reads were trimmed using Trimmomatic (v0.39) (Bolger et al. 2014) as described (Sielemann et al. 2022) and the quality was assessed using fastqc (v0.11.9) (Andrews 2020). All trimmed reads were subjected to DiscovarDeNovo (v52488) (run with default parameters and 10 threads; https://www.broadinstitute.org/software/discovar/blog) (Love et al. 2016) for *de novo* genome assembly after converting the fastq files to unmapped BAM files with picard tools (v2.5.0) (http://broadinstitute.github.io/picard/). Contigs with a length below 500 bp were discarded. BUSCO (v5.2.2) (Simão, Felipe A and Waterhouse, Robert M and Ioannidis, Panagiotis and Kriventseva, Evgenia V and Zdobnov, Evgeny M 2015) (embryophyta_odb10 dataset) was used with default parameters and 10 threads to assess the completeness of the assembly. Sequences with matches to a ‘black list’ were discarded as previously described (Siadjeu et al. 2020) to obtain the final genome assembly sequence (Supplemental_File_S6).

High-continuity long read assemblies were generated for *B. corolliflora*, *B. lomatogona*, and *P. procumbens*. Genomic DNA was extracted using a CTAB-based method (Siadjeu et al. 2020). Quality control was performed by agarose gel electrophoresis, NanoDrop measurement and Qubit analysis (Siadjeu et al. 2020). The short read eliminator kit (Circulomics) was used prior to library preparation following the SQK-LSK109 protocol. Sequencing was performed on a GridION using R9.4.1 flow cells as described previously (Siadjeu et al. 2020) (Supplemental_File_S5). For *B. corolliflora* and *B. lomatogona*, basecalling was performed using Guppy (v3.2) (https://nanoporetech.com/). Super high accuracy basecalling (v6) was available for read data from *P. procumbens*. A *de novo* assembly for each species was generated with Canu (v.1.8; for *P. procumbens*: v2.2) (parameters, excluding memory/threads: useGrid=1, saveReads=true, corMhapFilterThreshold=0.0000000002, ovlMerThreshold=500, corMhapOptions=--threshold 0.80, --num-hashes 512, --num-min-matches 3, --ordered-sketch-size 1000, --ordered-kmer-size 14, --min-olap-length 2000, --repeat-idf-scale 50) (Koren et al. 2017). Polishing of all assemblies was performed with racon (Vaser et al. 2017), followed by two rounds of medaka (https://github.com/nanoporetech/medaka) and three rounds of pilon (Walker et al. 2014) as described previously (Siadjeu et al. 2020). Contigs below 100 kb were discarded. ‘Decontamination’ of the assembly, i.e. discarding sequences with matches to a ’black list’, was performed as described previously (Siadjeu et al. 2020). The completeness of the final genome sequence assemblies (Supplemental_File_S6) was again assessed using BUSCO (v5.2.2) (Simão, Felipe A and Waterhouse, Robert M and Ioannidis, Panagiotis and Kriventseva, Evgenia V and Zdobnov, Evgeny M 2015) (embryophyta_odb10 dataset, -m genome, -c 10).

All assemblies were generated from DNA extracted from a single plant.

### Gene prediction and functional annotation

Prior to gene prediction, softmasking of the repeats in all genome assembly sequences was performed. A *de novo* repeat library was constructed with RepeatModeler (v2.0) (Flynn et al. 2020) including the LTR discovery pipeline. The RepBase library for each species together with the species-specific RepeatModeler library were used as input to RepeatMasker (v4.1.1) (Chen 2004).

The BRAKER2 pipeline (Brůna et al. 2021, 2; Lomsadze et al. 2014; Brůna et al. 2020; Lomsadze 2005; Buchfink et al. 2015; Gotoh 2008; Iwata and Gotoh 2012; Stanke et al. 2008, 2006) was used for gene prediction. Protein evidence, derived from OrthoDB protein sequences (Kriventseva et al. 2019) formatted with ProtHint (Brůna et al. 2020), as well as full-length sugar beet mRNA sequences from RefBeet-1.0 and RefBeet-1.5 were integrated as hints. The full-length mRNA sequences were aligned to the respective genome sequence assembly using BLAT (Kent 2002) (parameters: -fine; -q=rna). The alignments were filtered (filterPSL.pl; --best, --minCover=80, --minId=92), sorted by sequence names and begin coordinates, and then transformed into GFF format (blat2hints.pl). Both hint files, derived from RefBeet-1.0 and RefBeet1.5, were compared by alignment positions to discard the respective RefBeet-1.0 mRNA in case of an overlap with a RefBeet-1.5 mRNA. The alignments were merged to obtain the final hints file. The actual gene prediction was performed with BRAKER2 in the ‘etpmode’. Several scripts were used to reformat the resulting annotation file (fix_gtf_ids.py, gtf2gff.pl, augustus_to_GFF3_adapName.pl). Predicted genes with a resulting amino acid length below 50 were removed.

Each gene was named according to a species abbreviation composed of the first letter of the genus name and the first letter of the species name (e.g. *P. procumbens*: Pp). The contig name was added after an underscore and then followed by the gene number (sorted by assembly coordinates). The last part of the gene name is composed of four-letter codes - either based on reciprocal best hits (RBHs) identified by BLASTn against RefBeet genes, or by a new four-letter combination.

All genes were functionally annotated by InterProScan (v5.52) (Quevillon et al. 2005), SwissProt BLASTX (Altschul et al. 1990) and RBH-BLAST using published RefBeet annotations. The functional annotation files are available as part of this study (10.4119/unibi/2966932).

### Computational methods to resolve polyploid relationships

An overview of the read (Supplemental_File_S5) and assembly datasets (Supplemental_File_S6), used for the different approaches to resolve the parental relationships of *B. corolliflora*, is provided. Assembly statistics were calculated with QUAST (v. 5.2.0) (Gurevich et al. 2013).

### *K*-mer approaches to resolve polyploid relationships

To efficiently search specific *k*-mers in a given *k*-mer set, the *k*-mers were split into buckets based on the prefixes of length six. This corresponds to building a static trie (prefix tree) over the prefixes where the leaves of this trie are the buckets. As the trie already encodes the prefixes, only the suffixes in the buckets have to be saved in the form of sorted arrays. One can test for membership of a *k*-mer *m* by first traversing the trie along the prefix of *m* until a bucket is reached. If this traversal fails, *m* is not present in the set. If a bucket is reached successfully, a binary search in the bucket is performed. The application ‘*k*-mer operator’ (SBTTrio application; https://github.com/ksielemann/beet_pangenome) was written in Java.

To extract unique *k*-mers from a sequence, a sliding window of length *k* is moved over the sequence. All canonical *k*-mers (a canonical *k*-mer represents the lexicographically smaller of a *k*-mer and the corresponding reverse complement) are inserted into a hash table. The hash function is MurmurHash3 (Appleby). An open addressing hash table with a size that is always a power of 2 was used together with a quadratic probing function 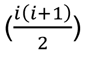 (Hopgood 1972).

The developed method can also be used to generate random sets of canonical *k*-mers. To achieve optimal time and space complexity at sampling (without replacement) random *k*-mers, a specific algorithm was used (sparse Fisher-Yates shuffle) (Ting 2021).

### - *K*-mer set operations

Similarities and differences between the species-specific *k*-mer sets were assessed using various set operations. As input for the *k*-mer set operations method, we first used normalised Illumina read datasets of the potential parent species to achieve a comparable set size. Normalisation was performed with bbnorm (Bushnell) and the parameters *k* = 21, a target depth of 20 and a minimum threshold of 3 to discard likely erroneous reads. For the child species, the complete set of *k*-mers based on the read datasets was used. In addition, the set operations were performed using sequence assemblies as input.

For each investigated species dataset, the set of distinct canonical *k*-mers (*k* = 13, 21, 31) was computed using the *k*-mer counting algorithm KMC3 (v3.2.1) (Kokot et al. 2017, 3). Based on the species-specific *k*-mer sets, multiple subsets were generated using different set operations. This includes the subset of *k*-mers present in all species (*B. corolliflora*, *B. macrorhiza*, and *B. lomatogona*), shared among two species as well as the subset of *k*-mers shared by two species, but not present in the third investigated species. These set operations were performed with KMC tools (v3.2.1) (Kokot et al. 2017, 3).

### **-** Generalised trio binning

Trio binning is traditionally used to separate reads into two haplotype-specific sets to generate phased assemblies (Koren et al. 2018). This approach was adapted to assess the parental contributions to tetraploid *B. corolliflora*. First, the *k*-mer set (*k*=21) for the potential parent species was calculated with KMC3 using either assemblies or high-quality, normalised (as described above) short reads as input. Then, the set of exclusive *k*-mers for each of the potential parent species was calculated with KMC tools (i.e. the set of *k*-mers present in one species, but not in the other). For the child species, a long-read dataset was used. For each read in this dataset, the number of unique canonical *k*-mers this read shares with the exclusive *k*-mer set of one of the parent candidate species was counted using the ‘*k*-mer operator’ described above. This number is then divided by the number of unique canonical *k*-mers of the read to get the ‘*k*-mer share’ for each potential parent. The average share of exclusive *k*-mers assigned to the potential parents was calculated over all reads. Only reads, for which at least half of the average *k*-mer share was assigned to the potential parents, are considered (the other reads contain too few *k*-mers exclusive to either one of the potential parental species). The goal of this filtering is to exclude reads where the overall signal is too weak to be interpreted reliably. All remaining reads were then classified into four different classes (Supplemental_File_S7): if the number of exclusive *k*-mers of a specific read is 3x higher for parent A than for parent B, the read was assigned to parent A i) (beige) and ii) *vice versa* (orange-red) (Supplemental_File_S7). The read was classified as ‘chimeric’ iii) if 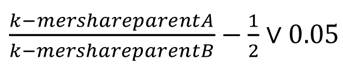 (orange). If none of the three conditions above applied, the read was ‘unclassified’ (IV) (grey). Generalised trio binning (SBTTrio application; https://github.com/ksielemann/beet_pangenome) was performed for the trio of interest (*B. corolliflora*, *B. lomatogona*, *B. macrorhiza*) as well as for other control trios to validate the approach (Table 2). *B. oleracea* (696 Mb) contributes a higher sequence content to allotetraploid *B. napus* in comparison to *B. rapa* (529 Mb) (Johnston 2005). For this reason, we normalised the results for this genome size difference.

### **-** *K*-mer fingerprinting

The *k*-mer fingerprinting approach introduces randomisation and is motivated by Fofanov *et. al.* (Fofanov et al. 2004) which suggests that small *k*-mer sets can be used to distinguish different organisms with high probability. The randomisation is also motivated by the prospect of minimising the impact of errors and therefore having a closer reflection of the real similarity when using the average over multiplerandom canonical *k*-mer sets. In general, this approach is similar to *k*-mer sketching.

To select a suitable *k*, the percentage of distinct canonical *k*-mers that are present in the dataset of each species was computed for a range of *k* (14-20). In accordance with Fofanov *et. al.* (Fofanov et al. 2004), a *k* was selected for which 5%-50% of all possible unique canonical *k*-mers were present in all datasets. This ensured that the different species datasets can be distinguished and that erroneous *k*-mers of low-quality reads do not impact the results. In general, the choice of *k* is a trade-off between a clear signal and computational intensity. In contrast to the previously described *k*-mer-based approaches, for which *k*=21 was well suitable, here, *k*=15 was selected based on the criterion by Fofanov *et. al.* 2004, and KMC3 was used to generate *k*-mer sets. The ‘*k*-mer operator’ was used to build indices for all investigated datasets and to generate 100,000 random sets of size 10,000 based on the whole set of all theoretically possible 15-mers. For each random set, the fingerprint, i.e. overlap with the species-specific *k*-mer set, was calculated. Additionally, the fingerprint intersection, i.e. the overlap of the fingerprint of the child species with the respective fingerprint of each investigated potential parent species, was computed.

*K*-mer fingerprinting was performed on assembly sequences for *B. corolliflora* as child species and *B. lomatogona*, *B. macrorhiza*, *B. patula*, *B. vulgaris* subsp. *maritima*, *B. vulgaris* subsp. *vulgaris*, and *P. procumbens* as candidate parent species.

### Mapping approach to resolve polyploid relationships

#### - Synthetic read mapping

As input, synthetic reads were generated from the sequence assembly of the child species (*B. corolliflora*). These synthetic reads were extracted by splitting each contig into equal length fragments starting from the beginning of the contig. The synthetic reads were then mapped simultaneously against the sequence assemblies of the two potential parent species (*B. lomatogona* and *B. macrorhiza*) using minimap2 within the corresponding Python wrapper mappy (v2.24) (Li 2018). The reads were then assigned to four categories either i) mapping to both potential parents, ii) mapping exclusively to parent A, iii) exclusively to parent B, or iv) not mapping to either of the parent candidates. A synthetic read length of 5 kb, 10 kb, and 20 kb was selected and only primary mappings with a sequence identity of at least 60% were considered. For comparability between the *B. lomatogona* long read-based and the *B. macrorhiza* short read assembly, in a second approach, the *B. lomatogona* assembly sequence was shredded into 5 kb (= smaller than the N50 of the *B. macrorhiza* assembly) chunks prior to the mapping procedure.

### Gene-based approach to resolve polyploid relationships

Phylogenetic distances of BUSCO gene sequences were calculated between tetraploid *B. corolliflora*, the potential parents, *B. vulgaris* subsp. *vulgaris*, *P. procumbens* and three related outgroup long-read genome sequence assemblies of the Caryophyllales (*Simmondsia chinensis* (jojoba; GCA_018398585.1), *Spinacia oleracea* (spinach; GCF_002007265.1), and *Amaranthus hypochondriacus* (amaranth; GCA_000753965.2)). For all eight species, BUSCO (v5.2.2) (Simão, Felipe A and Waterhouse, Robert M and Ioannidis, Panagiotis and Kriventseva, Evgenia V and Zdobnov, Evgeny M 2015) (embryophyta_odb10 dataset) was run in genome mode. Afterwards, suitable BUSCO genes were extracted. The final set of single copy (can be duplicated in the tetraploid *B. corolliflora*), complete BUSCO genes present in all six genome sequences comprised 140 genes. A multiple FASTA file was constructed for each gene and served as input for sequence alignment with MAFFT v7.299b (L-INS-I method; --adjustdirection) (Katoh, Kazutaka and Standley, Daron M 2013). The alignments were trimmed with trimAl (v1.4.rev22) (Capella-Gutierrez et al. 2009) to achieve 100% occupancy, which means that only SNVs were considered for the phylogenetic distance whereas InDels, possibly derived from assembly or gene structure annotation errors, were not considered. Single gene trees were constructed using FastTree (v2.1.11) (Price, Morgan N and Dehal, Paramvir S and Arkin, Adam P 2010). The phylogenetic distance of each parental gene to the closest related *B. corolliflora* gene was assessed using the DendroPy library (Sukumaran and Holder 2010). A Mann-Whitney-U test, implemented in the SciPy package (Jones et al. 2001), was calculated.

### Identification of lost/conserved regions in the pangenome

Illumina short reads of the sugar beet reference accession KWS2320 were used (Supplemental_File_S8) as input. After quality check, these reads were mapped against all six available wild beet genome sequence assemblies using BWA-MEM (v0.7.13) (-t 20, -c 1000) (Li 2013). In addition to the genomic resources presented in this study, we used two published genome sequence assemblies and annotations from *B. patula* and *B. vulgaris* subsp. *maritima* (http://bvseq.boku.ac.at/Genome/Download/) (Rodríguez del Río et al. 2019). The resulting SAM files were converted to BAM format using samtools (v1.15.1) (Li et al. 2009), then sorted (samtools sort), and duplicates were removed (samtools markdup). The whole workflow for the identification of regions of interest is visualised in Supplemental_File_S9, A. First, only mapped reads with a length greater than 80 bp and exclusively primary mappings were kept for further analyses to ensure a high-quality input dataset. The coverage per position was determined using genomeCoverageBed (v2.27.1) (-d, - split) (Quinlan and Hall 2010). Variant calling was performed with bcftools (v1.11) (Danecek et al. 2021) and the resulting variants were filtered for quality (QUAL Phred-score ≥ 30). Each position of the sequence was qualitatively assessed so that either a variant was present at a specific position (1), or no variant was detected (0).

The average coverage per base as well as the average variance per base was calculated in a sliding window approach. The approximate average gene length in sugar beet is 5 kb (based on the KWS2320ONT v1.0 p1.0 annotation). To ensure high sensitivity and as consecutive conserved regions are later merged into a single window, a window size of 2,500 bp was selected. An example region is shown in Supplemental_File_S10. The shift size of 150 bp means that the first window spans the region from 0 bp to 5,000 bp, whereas the second window spans the region from 150 bp to 5,150 bp (Supplemental_File_S9, B). All parameters can be selected by the user depending on the application.

As stated above, regions not present in cultivated sugar beet KWS2320 should be associated with low/no coverage in the crop wild relatives. To get these ‘zero coverage regions’, after the extraction of primary mappings, a maximal coverage of 1% of the mean coverage per contig was set as filter criterion. As short-read assemblies are highly fragmented and short contigs are present, the coverage was normalised in these cases by the value calculated from the whole assembly (for *B. vulgaris* subsp. *maritima*, *B. patula*, and *B. macrorhiza*).

As *B. vulgaris* subsp. *maritima* is known to be the progenitor of the cultivated sugar beet (Biancardi and Lewellen 2020), the pangenome dataset was also harnessed to identify regions originally derived from the progenitor (sea beet WB42) and still conserved in the descendant (cultivated sugar beet KWS2320) (Supplemental_File_S4). A conserved region between sea beet and sugar beet was defined as follows. If the region derives from sea beet, the *B. vulgaris* subsp. *maritima* reads should map to the corresponding, conserved region in the sugar beet genome sequence. Therefore, the coverage should be at least as high as the mean coverage across the respective contig. To exclude highly repetitive sequences, an upper threshold was defined as well (at most 3x the mean coverage for each contig). On the other hand, the expected variance for conserved and therefore similar regions in both sequences should be relatively low. The maximal variance per base threshold was set to 0.4 times the average variance per base of the respective contig (sum of all variants of the contig divided by the contig length x 0.4).

All identified lost/conserved regions were then further investigated. First, based on the structural annotations of all assemblies, gene sequences, which are located within the identified regions, were extracted. Genes were considered to be located within a specific region if at least 70% of the bases were covered. The corresponding amino acid sequences were used for the next step. To functionally characterise the extracted genes, RBHs with *A. thaliana* amino acid sequences were determined and the corresponding *A. thaliana* gene identifiers were assigned to the respective beet gene. In addition, the functional annotation file for *B. corolliflora*, *B. lomatogona*, *B. macrorhiza*, and *P. procumbens*, which was generated in this study (10.4119/unibi/2966932), was investigated to further functionally assess the identified genes.

### Chromosome preparation and fluorescent *in situ* hybridisation

Mitotic chromosomes were prepared from young meristematic leaves of *B. corolliflora* (BETA 408), *B. lomatogona* (BETA 674) and *B. macrorhiza* (BETA 830) as described previously (Schmidt et al. 2021, 2023). Probes for the satellite DNAs pRN1 (Kubis et al. 1997), GenBank accession number Z69354.1) and BlSat1 (Hong Ha 2018) were labelled by PCR in the presence of biotin-16-dUTP (Roche Diagnostics) detected by streptavidin-Cy3 (Sigma–Aldrich) or digoxigenin-11-dUTP (Jena Bioscience) detected by antidigoxigenin-fluorescein isothiocyanate (FITC; Roche Diagnostics). The 18S rDNA probe was labelled with DY415-dUTP (Dyomics). All probe nucleotide sequences are listed in the Supplemental_File_S11. Chromosomes were counterstained with DAPI (4′,6′-diamidino-2-phenylindole; Böhringer, Mannheim) and mounted in antifade solution (CitiFluor; Agar Scientific, Stansted). The hybridization procedure as well as the image acquisition were performed as described previously (Schmidt et al. 2021; Liedtke et al. 2022). The hybridization stringency was 79%.

## Declarations

### Ethics approval and consent to participate

The material of the IPK Gatersleben was transferred under the regulations of the standard material transfer agreement (SMTA) of the International Treaty. Plants were grown in accordance with German legislation.

### Consent for publication

Not applicable.

### Availability of data and materials

ONT reads, Illumina reads, and genome assemblies generated for this study were submitted to ENA (PRJEB56520). The sources/IDs are summarised in Additional files S5 and S6. The structural and functional annotation files for all generated wild beet assemblies are available on ‘PUB-Publications at Bielefeld University’ (10.4119/unibi/2966932). Relevant scripts for the investigation of the parental relationships of *B. corolliflora* and for the identification of lost/conserved regions are available on GitHub (https://github.com/ksielemann/beet_pangenome; https://doi.org/10.5281/zenodo.8090593).

### Competing interests

The authors declare no competing interests.

## Funding

KS is funded by Bielefeld University through the Graduate School DILS (Digital Infrastructure for the Life Sciences). The bench fee was contributed from internal resources of the chair of Genetics and Genomics of Plants through core funding from Bielefeld University/Faculty of Biology.

### Authors’ contributions

KS, BP, BW, TH, and DH designed the study. NS selected and cultivated the plants. NS, KS, and BP performed DNA extraction. PV and BP designed the layout for sequencing and performed sequencing. KS, JG, and NK developed and implemented the bioinformatic methodology. SB and NS performed the generation of probes and hybridisation experiments. KS, NS, JG, and NK analysed the data and prepared the figures and tables. KS, NS, JG, NK, BP, and TH wrote the manuscript. All authors read and approved the final manuscript.

## Acknowledgements

We thank the CeBiTec Bioinformatic Resource Facility team for excellent technical support. We acknowledge the Genbank Gatersleben for providing seeds and data for the investigated accessions. This work was supported by the BMBF-funded de.NBI Cloud within the German Network for Bioinformatics Infrastructure (de.NBI) (031A532B, 031A533A, 031A533B, 031A534A, 031A535A, 031A537A, 031A537B, 031A537C, 031A537D, 031A538A).

## Notes

### Competing Interest Statement

The authors have declared no competing interest.

